# High throughput identification of genetic regulators of microglial inflammatory processes in Alzheimer’s disease

**DOI:** 10.1101/2025.03.09.642133

**Authors:** Christopher L. Cardona, Lai Wei, Joonwon Kim, Ellen Angeles, Gunjandeep Singh, Shiye Chen, Ronak Patel, Nkechime Ifediora, Peter Canoll, Andrew F. Teich, Gunnar Hargus, Alejandro Chavez, Andrew A. Sproul

**Author notes:** Denotes equal authorship contributions (Christopher L. Cardona, Lai Wei, Joonwon Kim, Ellen Angeles). Corresponding Author: Alejandro Chavez, Lead Contact Andrew A. **Data availability statement**: All data used in this study are presented in the main text and/or the Supplementary Materials. The raw sequencing data from our CRISPRi screening experiments have been deposited in the NCBI Sequence Read Archive (SRA) under BioProject accession number PRJNA1152984. All raw single-cell RNA sequencing data have been submitted to GEO accession # GSE289721. **Conflict of interest statement:** A.C., J.K., and L.W. have patents related to CRISPR-based tools and their application that are managed by Harvard and Columbia University.

## Abstract

Genome-wide association studies (GWAS) have identified over a hundred genetic risk factors for Alzheimer’s disease (AD), many of which are predominantly expressed in microglia. However, the pathogenic role for most of them remains unclear. To systematically investigate how AD GWAS variants influence human microglial inflammatory responses, we conducted CRISPR inhibition (CRISPRi) screens targeting 119 AD GWAS hits in hiPSC-derived microglia (iMGLs) and used the production of reactive oxygen species (ROS) in response to the viral mimic poly(I:C) as a functional readout. Top hits whose knockdown either increased or decreased ROS levels in response to poly(I:C) were further analyzed using CROP-seq to integrate CRISPRi with single-cell RNA sequencing (scRNA-seq). This analysis identified 9 unique microglial clusters, including a poly(I:C)-driven inflammatory cluster 2. Emerging evidence supports a pathogenic role of viral infections in AD and cross comparison of our scRNA-seq data with iMGLs xenotransplanted into an AD mouse model shows significant overlap between our clusters and AD-relevant microglial clusters. Knockdown of *MS4A6A* and *EED*, which resulted in elevated ROS production in the presence of poly(I:C), increased the proportion of cluster 2 cells and induced functionally related changes in gene expression. In addition, KD of *MS4A6* led to a reduction in the proportion of iMGLs in the DAM (disease associated microglia) cluster under all conditions, suggesting that this gene may modulate the DAM response. In contrast, KD of *INPP5D* or *RAPEP1* which lead to low levels of ROS in the presence of poly(I:C), did not significantly affect the proportion of cells in cluster 2 but rather shaped the inflammatory response. This included the upregulation of an HLA-associated inflammatory cluster (cluster 6) by *INPP5D* knockdown under all conditions, independent of poly(I:C) stimulation. Importantly, KD of *INPP5D* or *RAPEP1* had many shared differentially expressed genes (DEGs) under both vehicle and poly(I:C) treated conditions. Overall, our findings demonstrate that despite the diverse biological functions of AD GWAS variants, they converge functionally to regulate human microglial states and shape inflammatory responses relevant to AD pathology.

## 1 Introduction

Large-scale genome-wide association studies (GWAS) have implicated greater than 100 genes in the pathogenesis of Alzheimer’s disease (AD), yet functional proof of their significance using experimental models is lacking for the vast majority. In addition, it has become apparent that human cells react differently to AD pathology than their murine counterparts. For example, human hiPSC-derived microglia (iMGLs) xenotransplanted into a humanized AD mouse model (*APP^NL-G-F^*) have cytokine response microglia (CRM) populations not seen in resident mouse glia^1^. Human versions of some AD-related genes also have different properties than murine versions, such as tau isoform composition^2^. Thus, looking at AD risk genes in human induced pluripotent stem cell (hiPSC)-derived models provides a unique opportunity to better understand the drivers of AD pathogenesis, which can be explored at larger scales by using CRISPR screening approaches^3–6^.

For these studies we have focused on microglia given an ever-increasing understanding of the critical roles they play within both homeostatic and disease context. Microglia are brain resident immune cells that provide trophic support to neurons and promote neuronal plasticity, synaptic pruning and help maintain brain homeostasis. In response to stimuli microglia can develop into neuroprotective or pro-inflammatory subtypes with expression of anti-or pro-inflammatory molecules, respectively^7^. Microglia are activated under various conditions including trauma, ischemia, infection, inflammation, and neurodegeneration^7^. As such, imaging studies in AD patients have demonstrated that microglial activation begins early during disease development, and that their activation appears to be highest in areas of neuronal cell death, where they may play critical roles in removing apoptotic cells from the brain and contribute to disease pathogenesis^7^. Recent single cell studies also indicate the presence of additional microglial subpopulations in AD brains with interferon-related, proliferation-related, or lipopolysaccharide-related transcriptome signatures, which may have different functions in driving the underlying disease pathobiology^7–9^. Importantly, many AD genetic risk factors are primarily expressed in microglia^10,11^ suggesting it is particularly critical to assess the function of AD GWAS risk factors in this cell type. Thus, we developed a high throughput CRISPRi screening approach to look at many AD GWAS risk factors in parallel for effects on microglia response to inflammatory stimuli, followed by single cell analysis of our top hits to better understand their effects on microglial cell states and inflammatory signalling. This led to the identification of genes that drove inflammatory processes when knocked down and genes that when knocked down modulated inflammatory processes and significantly affected the microglial transcriptome under steady-state conditions.

## 2 Methods

### 2.1 Cell culture and differentiations

The FA0000010 (FA10; RUCDR/Infinity BiologiX) hiPSC line used in this study was maintained feeder free in StemFlex (ThermoFisher) on Cultrex (ThermoFisher) and split 1-2 times a week with ReLeSR (STEMCELL Technologies) as has been described previously^12^. Differentiations into human microglia (induced microglia-like cells; iMGLs) was done using a previously published protocol^13^ with minor modifications as we have done previously^14–16^ (Figure S1), which includes generating bankable hematopoietic progenitors (HPCs) using a commercial kit (STEMCELL Technologies). Human embryonic kidney 293 (HEK293) cells were grown in 10% FBS/DMEM (ThermoFisher) as described previously^12^.

### 2.2 Plasmid construction and lentiviral preparations

The gRNA sequences in our CRISPRi library were primarily adapted from the well-cited Dolcetto library from the Broad Institute^3^. Additional guides were designed for genes that were not present in library using Broad’s Genetic Perturbation Platform sgRNA designer^4^. Each gene in the final libraries was represented by 6 guides. Our oligo design included two BSMBI sites to allow for Golden Gate Assembly into our desired backbone. Our 119-base pair oligonucleotides with spacer sequences (**Table S1**) were synthesized on an oligochip (Agilent Technologies). The gRNAs were then amplified from the oligopool and cloned into the pSB700-puro F+E vector (Addgene: #167927) using Esp3I, as described previously^17^. The gRNA library was then transfected as a mixed pool into HEK293 cells to generate lentivirus for transducing microglia. Lentiviruses was concentrated with Lenti-X concentrator (Takara) as has been done previously^12^.

For CROP-seq analysis, we generated a modified the CROP-seq vector (based on Addgene: # 106280) carrying a PuroR-T2A-mVenus selection marker along with incorporating the 10x Genomics capture sequence (5’-GCTCACCTATTAGCGGCTAAGG-3’) at the 3’-end of the gRNA scaffold. Three distinct gRNAs per gene, selected from our previous screen (**Table S2**), and were individually cloned into this modified vector as previously described^18^. Midipreps (Macherey-Nagel) were conducted individually for all 24 gRNA constructs and 3 gRNAs for each gene were combined to make one 1 lentiviral preparation per gene.

### 2.3 Generation of FA10 dCas9-KRAB-MeCP2 lines

FA10 hiPSCs were infected with dCas9-KRAB-MeCP2^19^ (Addgene: #122205) lentiviral particles by reverse transduction and selected with 5 μg/mL blasticidin to generate permanent polyclonal lines, as has been done in a similar fashion previously for other transgenes^12^. Two independent clones were generated and validated for KD using a gRNA against the well-expressed cell surface protein EPCAM. For all studies clone Cl.1 was used as it showed better KD within iMGLs in pilot experiments.

### 2.4 Pooled CRISPRi screens in iMGLs

A pooled gRNA library containing 1026 guide RNAs targeting 171 genes was delivered via lentiviral transduction into the dCas9-KRAB-MeCP2 overexpressing iMGLs at day 21 of differentiation post HPC at a low multiplicity of infection (MOI) of approximately 0.3, which was determined by pilot experiments utilizing empty GFP virus. This low MOI was chosen to ensure that most transduced cells received at most one gRNA, thus generating a pool of microglia with primarily a single gene perturbed. The transduced iMGL pool was treated at with 10 μg/mL Poly(I:C) at day 35 of differentiation post HPC to stimulate an inflammatory response. Following a 2h treatment, cells were stained with 1 μM CellROX Deep Red (ThermoFisher) to measure the intracellular levels of reactive oxygen species (ROS), as per the manufacturer’s instructions. Stained cells were then sorted using Fluorescence-Activated Cell Sorting (FACS) based on CellROX intensity (Columbia Stem Cell Initiative Flow Cytometry Core). Four populations were collected: Top 12.5% CellROX intensity (T1), Top 12.5-25% CellROX intensity (T2), Bottom 12.5-25% CellROX intensity (B2), Bottom 12.5% CellROX intensity (B1). For each sorted population, ∼500,000 cells were collected, pelleted, and frozen. Genomic DNA from sorted iMGLs was extracted by adding 50 μL of Lucigen QuickExtract solution, followed by heating the sample according to the manufacturer’s instructions. The gRNA distribution within cells was analyzed via next generation sequencing as previously described^17^. Two independent iMGL differentiations and subsequent CRISPRi screens were conducted.

### 2.5 Data Preprocessing for CRISPRi screens

Raw sequencing data from the pooled CRISPRi screen were processed by MAGeCK (Model-based Analysis of Genome-wide CRISPR-Cas9 Knockout) software package (version 0.5.9)^20^ to obtain read counts for each gRNA. Quality control metrics, including total read count and gRNA distribution were assessed to ensure data reliability. Read counts were normalized using the median normalization method implemented in MAGeCK to account for differences in sequencing depth between samples. As an initial assessment of CRISPRi activity we calculated overall depletion of essential gRNAs within the library using the MAGeCK test command comparing gRNA abundance within the pooled microglia and the plasmid library used for lentivirus packaging (**Figure S1**; **Table S3**). This step calculates a score and p-value for each gRNA based on its change in abundance. Gene-level significance was determined using the robust ranking aggregation (RRA) algorithm implemented in MAGeCK. This approach combines the effects of multiple gRNAs targeting the same gene to identify significantly enriched or depleted genes. Two separate CRISPRi screens were analyzed using the MAGeCK-RRA algorithm to produce two independent ranked lists of genes exhibiting either depletion or enrichment during treatment. Genes common to both lists in the top 20 for either T1/B1 or B1/T1 were selected as the screen hits for further investigation.

### 2.6 CROP-seq experiments

Two independent CROP-seq experiments were conducted^21^. For each experiment, 2 independent 12-well wells of cells differentiated 21 days post HPC were infected with 3 gRNAs corresponding to one of 8 target genes (High ROS: *AGF2*, *EED*, *MS4A6*; low ROS: *INPP5D*, *PVR*, *RABEP1*; negative controls (CTRL): *CABP5*, *SAGE1*). On day 35, one well for each target gene was treated with vehicle or 10 μg/mL poly(I:C) for 2h, dissociated with Accutase, pooled into either vehicle or poly(I:C) treatment Eppendorf tubes at ∼1000 cells/μL and processed for single cell RNA sequencing (scRNA-seq)^22^.

### 2.7 CROP-seq analysis

scRNA-seq was performed at the JP Sulzberger Columbia Genome Center single-cell sequencing core on a 10x Genomics Chromium Single Cell Controller. For each sample, a library was prepared for gene expression analysis along with a secondary library enriched for the gRNAs within the population of cells. Raw FASTQ files were aligned to the human transcriptome (version refdata-gex-GRCh38-2020-A, 10x Genomics) and transcripts were quantified using cellranger count (version 7.1.0, 10x Genomics) with additional parameters for CRISPR guide capture. Subsequent analysis was performed in R (version 4.2.2). Filtered count matrices were imported into R and converted to SeuratObjects. Quality control was performed by eliminating cells that: (i) had less than 750 or greater than 6,680 number of genes detected per cell; (ii) had less than 500 or greater than 28,500 number of unique molecular identifiers per cell; and (iii) had greater than 12% mitochondrial gene expression (genes present in mitochondrial DNA). We performed integration using canonical correlation analysis to remove batch effects from different experimental replicates. Then, we used Seurat^23^ (version 5.0.2) to perform normalization, identification of highly variable features, scaling, linear dimensionality reduction, Louvain clustering, and non-linear dimensionality reduction using UMAP. We used clustree^24^ (version 0.5.1) to identify the optimal number of clusters over multiple resolutions.

#### CROP-seq analysis with Mixscape

We used the workflow described for Mixscape within Seurat^25^. To assign perturbations to cells, we used the CRIPSR guide counts to define the expected perturbation within each cell. Non-targeted cells were defined as having less than 5 counts for all gRNA sequences. For all other cells, we assigned the perturbation based on the guide that had the highest number of counts for that cell. We segregated cells based on treatment and performed Mixscape on vehicle and poly(I:C)-treated iMGLs to classify perturbed cells.

#### Proportion analysis

Differences in cluster proportions between vehicle and poly(I:C)-treated iMGLs and between CRISPRi targets and CTRL were analyzed using scProportionTest (version 0.0.0.9)^26^ with default parameters. We highlighted clusters that were significantly depleted or enriched (observed log2 fold difference ≥ 0.5). Significance was set according to FDR ≤ 0.05.

#### Cluster markers and differential expression

Cluster markers were identified using FindMarkers in Seurat with default parameters. We selected only those significantly upregulated genes with an average log2 fold change ≥ 0.5. DEGs between vehicle and poly(I:C)-treated iMGLs were identified using FindMarkers with default parameters. We selected only those significantly upregulated genes with an average log2 fold change ≥ 1. We identified target specific DEGs by comparing CRISPRi classified cells for a particular target against CTRL. ggVenn (version 0.1.10) was used to compare DEGs between targets. Significance was set according to adjusted *p*-value ≤ 0.05.

#### Pathway analysis

DEGs were used as input for pathway analysis using EnrichR^27^ (version 3.2). GO biological processes 2023 database was used to identify pathways associated with significant DEGs. For each pathway, the gene ratio was calculated by dividing the number of input DEGs that were part of a given pathway by the total number of genes in that pathway. Top significant pathways were used for visualization. Significance was set according to adjusted *p*-value ≤ 0.05.

#### Comparison of clusters with xenotransplanted microglia

To assess how similar our iMGL clusters corresponded with other scRNA-seq data from xenotransplanted iMGLs (xMGLs)^1^, we used Seurat’s AddModuleScore function. This was used to assess the average expression of a set of genes (all iMGL clusters) from Mancuso et al. among our iMGL clusters. As input, we selected the top significant markers from each xMGL cluster with a log fold change greater than 0.5. Raw module scores were converted to z-scores by comparing the cluster means to the global mean across all clusters. A two-tailed p-value was derived from the z-scores to assess significant changes in module scores from the global mean. Raw p-values were adjusted for multiple comparisons using the Bonferroni method.

## 3 Results

### 3.1 Generation of pooled CRISPRi gRNA libraries for AD GWAS hits and control genes

Two large-scale AD GWAS studies^28,29^ were used to generate 119 targets for our CRISPRi gRNA library (**Figure 1A**). 49 genes (Kunkle et. al) were chosen because they have been identified as containing or being proximate to late onset AD risk loci by GWAS^1^. These genes were prioritized based on comparisons to other AD studies including loss-of-function assays and differential expression analysis between AD brain samples compared to control samples. The remaining 70 genes of interest (Jansen et al.) have been identified as potentially important in AD pathogenesis through a GWAS focusing on both diagnosed AD patients as well as AD-by-proxy (direct family members of diagnosed AD patients)^2^. Genes were identified from identified risk loci through a combination of positional gene-mapping, expression quantitative trait loci (eQTL) mapping, chromatin interaction mapping, and genome-wide gene-based association analysis (GWGAS) statistical mapping^2^. Two additional sets of control genes were also included. 26 essential genes as assessed by DepMap^30^, a data repository with over 1,000 CRISPR screens, were included as positive controls as they should be depleted by successful CRISPRi. 26 non-essential genes as determined by DepMap were included as negative controls that would not be predicted to be depleted.

**Figure 1:**
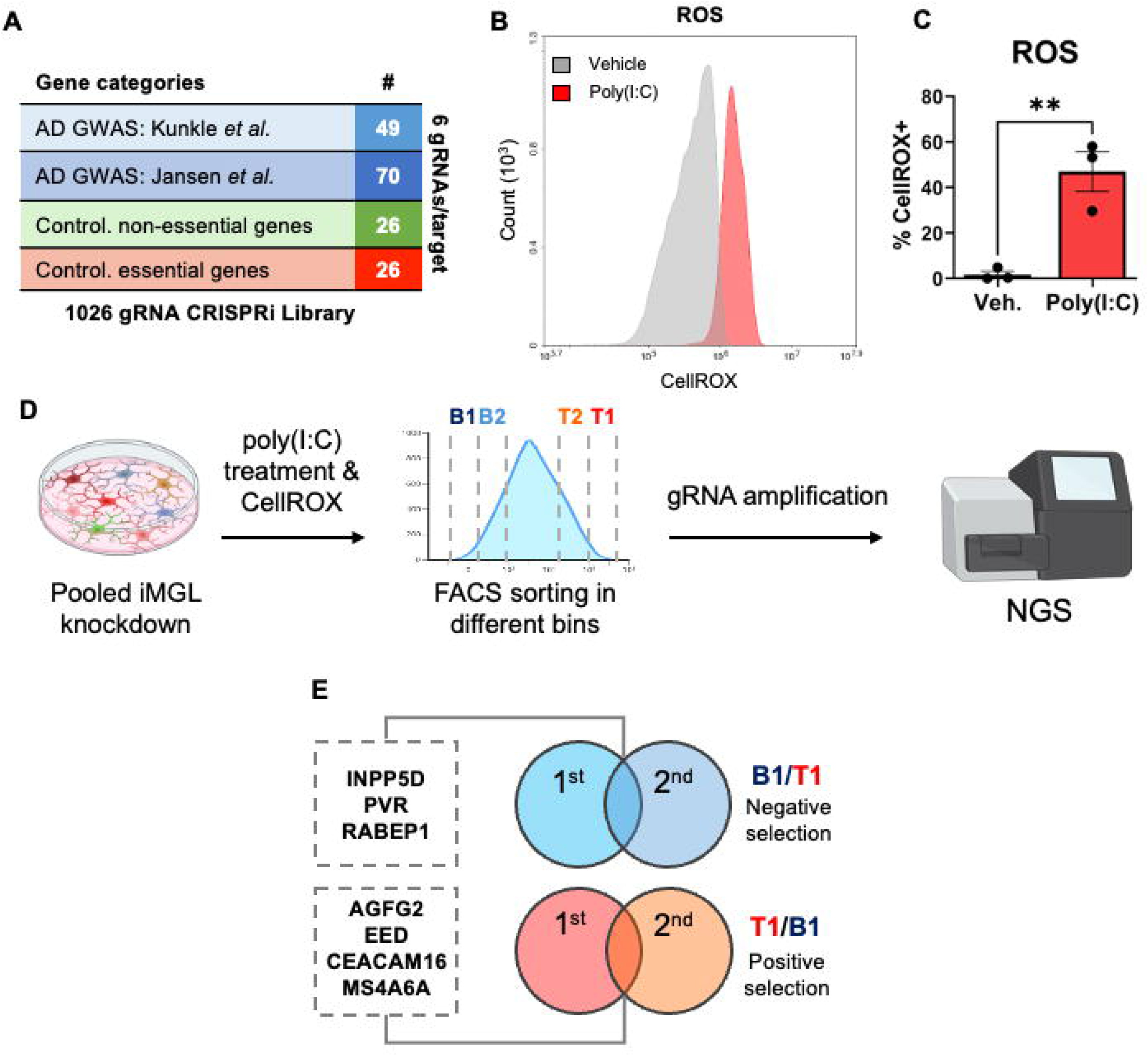
CRISPRi screens of AD GWAS hits characterizing their effect on ROS production. (**A**) Number of targets for 4 categories of genes for the CRISPRi sgRNA library. 6 sgRNAs were utilized per gene, for a total of 1026 gRNAs. (**B-C**) Treatment with poly(I:C) for 2h led to an increase of ROS as measured by flow cytometry (FC) of CellROX+ cells. Representative FC histogram is shown in (**B**) and quantification in (**C**). ** = p<0.01, Student’s t-Test, N=3 biological replicates, error bars are S.E.M. (D) Cartoon schematic of large-scale CRISPRi screens done in 2 independent differentiations. (**E**) Genes consistently in the top 20 for negative selection (B1/T1 ratio greater) or positive selection (T1/B1 ratio greater) for 2 independent screens as measured by the MAGeCK RRA algorithm are shown.

### 3.2 Generation of dCas9-KRAB-MeCP2 hiPSCs and development of a fluorescent readout for NGS

After our CRISPRi gRNA library was in hand we needed to generate an efficient method of knocking down target genes. To do so, we took the FA0000010 (FA10) cell line and introduced a dCas-KRAB-MeCP2^19^ cassette by lentiviral infection and subsequent blasticidin selection. FA10 was selected because we previously found this line robustly differentiates into iMGLs. Two independent polyclonal hiPSC lines were generated and validated for CRISPRi efficacy by using a previously validated gRNA targeting EPCAM (data not shown).

We next developed an assay that could measure inflammatory responses through an easy to quantify fluorescent readout. We chose to use reactive oxygen species (ROS) production as measured by CellROX (Deep Red) in response to the inflammatory stimuli poly(I:C), a double-stranded RNA (dsRNA) viral mimic. It is becoming increasingly evident that viral and other microbial infections likely play pathogenic roles in AD^31,32^, and it is important to interrogate how AD GWAS factors affect microglial responses to these stimuli. FA10 hiPSCs were differentiated into iMGLs through bankable hematopoietic progenitors (HPCs)^13^, as we have done previously^14–16^. Treatment of iMGLs led to an increase in CellROX fluorescence intensity (**Figure 1B-C**) which we then could use as a readout for our CRISPRi screens.

### 3.3 CRISPRi screens of AD GWAS hits for regulators of ROS production

Our CRISPRi screening strategy is shown schematically in **Figure 1D**. We conducted pilot transduction experiments at differentiation day 21 post HPC to experimentally derive a ∼30% infection rate at day 35 so that each iMGL would only be expected on average to receive 1 gRNA from our CRISPRi library (data not shown). We then carried out two independent large-scale CRISPRi screens in iMGLs treated with poly(I:C), where the top and bottom quartile of CellROX expression was subjected to fluorescence-activated cell sorting (FACS) into 2 bins each. Next generation sequencing (NGS) was conducted and genes that were consistently enriched or depleted for CellROX staining within two independent screens were then further analyzed (**Figure 1E; Table S3**). Importantly, positive control essential genes were depleted in each screen as expected (**Figure S1**). We refer to “high ROS hits” as genes that increased ROS levels in response to poly(I:C) when knocked down and “low ROS hits” as genes that lowered ROS in response to poly(I:C) when knocked down.

### 3.4 CROP-seq investigation of top AD GWAS ROS modulators

Three high ROS hits (*AGF2*, *EED*, *MS4A6A*) and three low ROS hits (*INPP5D*, *PVR, RABEP1*) were further analyzed in two independent CROP-seq experiments. Two non-essential genes (*CABP5*, *SAGE1*) that are not expressed in iMGLs (unpublished bulk RNA-seq data) were used as negative controls. Differentiating iMGLs were transduced with gRNAs against each gene of interest on differentiation day 21 post HPC, and half the cells were treated with vehicle and half with poly(I:C) for 2h on day 35 post HPC and subjected to scRNA-seq. 4 of 6 genes tested had significant effects on gene expression (*INPP5D*, *RABEP1*, *MSA4A6A*, *EED*) and were analyzed further.

### 3.5 Analysis of the effect of poly(I:C) on microglial cell state

When incorporating all iMGLs from our 2 independent screens, we identified 9 total microglial clusters in our dataset (**Figure 2A**) and observed that iMGLs were enriched to specific clusters based on treatment conditions (**Figure 2B**). When we looked at the distribution of cells across clusters for vehicle and poly(I:C)-treated iMGLs, we noticed differences in cluster percentages (**Figure 2C; Table S4**). Most notably, we saw a dramatic increase in cluster 2 in poly(I:C)-treated cells, having 34.2% of cells in this cluster compared to 1.2% of cells from vehicle-treated cells. We performed proportion analysis to assess which clusters were significantly depleted or enriched in poly(I:C)-treated iMGLs. We observed a highly significant enrichment of cells in cluster 2 for poly(I:C)-treated cells, with a log2 fold difference greater than 4. We also observed significant depletion of cells in clusters 0, 3, and 5, with log2 fold differences near 1 (**Figure 2D**). The maintenance of all clusters even in the setting of a highly inflammatory stimulus highlights the heterogeneity in microglia responses and suggests that a complex interplay between various microglia cell states may underlie their role within *in vivo* settings such as the inflammatory milieu present within LOAD.

**Figure 2:**
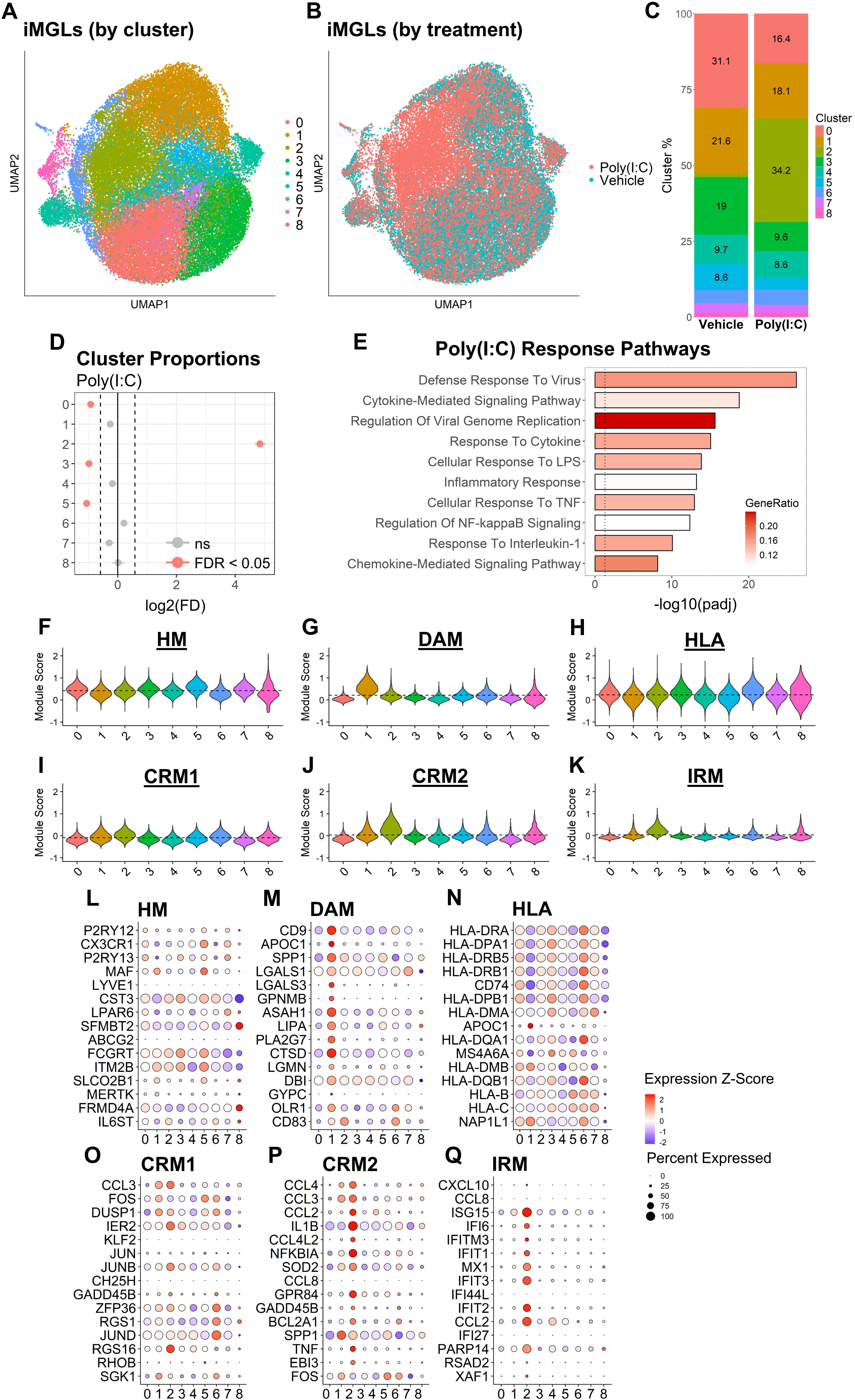
Characterization of iMGL responses to poly(I:C). (**A**) UMAP visualization of 44,913 single cells from iMGLs treated with either vehicle or the viral mimic poly(I:C) and colored according to 9 identified microglial clusters or (**B**) by treatment. (**C**) Cluster percentages for vehicle and poly(I:C)-treated iMGLs. (**D**) Point-range plot showing observed log_2_ fold difference for each cluster with dotted lines indicating a minimum log_2_ fold difference of 0.5. Points colored red are significant (FDR <0.05). (D) Pathway analysis for 182 significant DEGs comparing poly(I:C) to vehicle with a log_2_ fold change of at least 1. The top 10 significant GO biological process pathways are colored according to gene ratio. (**F-K**) Violin plots showing the distribution of module scores across clusters for homeostatic (HM, F), DAM (G), HLA (H), cytokine-response microglia 1 (CRM1, I), cytokine-response microglia 2 (CRM2, J), and interferon-response microglia (IRM, K) based on data from xenotransplanted iMGLs (xMGLs)^1^. The dotted line indicates the global mean of module scores across all clusters. (**L-Q**) For each module from xMGL dataset, the dot plots show how the top 15 markers from that dataset are expressed in our iMGL clusters. The color of the dots represents the expression z-score across all cells in that cluster and the size of the dot represents the percentage of cells in which the gene was expressed.

We then wanted to know which pathways were associated with DEGs for poly(I:C)-treated iMGLs relative to vehicle-treated cells. Pathway analysis uncovered key functional pathways within GO biological processes. We identified 595 significant pathways associated with 182 upregulated (UR) DEGs in response to poly(I:C) (**Table S4**). The top pathways were related to antiviral responses, response to cytokines and chemokines, inflammatory responses, NFκB signaling, and TNF signaling (**Figure 2E**). To further characterize human iMGL states, we compared our clusters with those from another single-cell RNA dataset of xenotransplanted iMGLs (xMGLs) into the humanized amyloid mouse model (*APP^NL-G-F^*). We selected the top significant markers across their homeostatic (HM), DAM, HLA, cytokine-response microglia 1 (CRM1), cytokine-response microglia 2 (CRM2), and interferon response microglia (IRM) clusters with a log2 fold change greater than 0.5 to use as genes which defined those clusters. We used those markers to calculate a module score for each of our clusters, which indicates the average expression of that gene set in each cell. We found that clusters 0, 3, and 5 had above average module scores for the previously defined homeostatic microglia cluster, with higher expression of key homeostatic genes, such as *CX3CR1* and *P2RY12*, compared to other clusters (**Figure 2F,L, Figure S2,** and **Table S4**). Additionally, we observed strong depletion of these clusters in response to poly(I:C), as compared to other clusters. Taken together, we propose that clusters 0, 3, and 5 may include homeostatic microglia, which represent a cellular reserve that is better primed to rapidly change transcriptional state versus other microglia populations in the setting of an acute inflammatory stimulus. Module scores for xMGL DAM genes were drastically increased in cluster 1, with a set of clearly defined marker genes that were uniquely expressed in that cluster (i.e., *PLA2G7*, *TREM2*, *GPNMB*, *MYO1E*, *NPL*) and concordant with known markers from DAM xMGLs and other reported human DAM clusters^33–35^ (**Figure 2G,M** and **Figure S2**). The cells in cluster 1 were present in both control and poly(I:C) conditions, with only a slight change in their abundance upon exposure to a strong acute stimulus. These data suggest that this population is not unique to disease states and may represent one of several stable subpopulations that are not rapidly altered upon some acute inflammatory stimuli. There were numerous significant pathways associated with marker genes in this cluster, including lipid metabolism, iron homeostasis, and receptor-mediated endocytosis pathways (**Table S4).** For the HLA module, we found that cluster 6 had a significantly higher module score compared to other clusters, with a module score z-score of 35.7, and had the highest expression of HLA markers expressed in xMGLs (**Figure 2H,N** and **Table S4**). Cluster 6 exhibited activation of numerous pathways associated with immune responses and included response to cytokines, regulation of p38MAPK signaling, and NF-κB signaling (**Figure S2** and **Table S4**). As HLA cluster 6 is not upregulated by poly(I:C), it suggests a narrower inflammatory role for this population. HLA+ antigen-presenting microglia, including xMGLs, have been identified in close proximity with amyloid plaques^1,36^. Lastly, we observed the cluster 2 had the highest module score z-scores associated with CRM1, CRM2, and IRM clusters from xMGLs with high expression of pro-inflammatory and antiviral genes (*IFIT2*, *IFIT3*, *MX1*, *IL1A*, *RELB*) (**Figure 2I-K,O-Q, Figure S2,** and **Table S4**). There were many significant pathways associated with genes in this cluster, which included pathways associated with viral defense responses, inflammation, and regulation of NF-κB signaling (**Table S4**). We also identified 2 clusters associated with proliferation, 4 and 7, which had overlap of some marker genes (**Figure S2**). Associated pathways in these clusters were related to DNA replication, DNA repair, and mitotic spindle organization. Lastly, while cluster 8 was small, there was a clear set of markers genes with many significant associated pathways. These were related to chromatin remodeling, epigenetic regulation, and vesicle-mediated transport (**Figure S2** and **Table S4**). The small numbers and associated pathways within cluster 8, suggest that it may represent some transient cellular state undergoing active rewiring, although future studies will be required to confirm this finding.

### 3.6 KD of ROS modulators change microglial cell states

Next, we wanted to understand how CRISPRi-mediated knockdown of our AD GWAS hits that modulate ROS affected microglial cluster proportions. Each target gene from low (*INPP5D*, *RABEP1*) and high (*MS4A6A*, *EED*) ROS populations were compared against negative control gRNAs targeted to non-essential genes which are not expressed in iMGLs (*CABP5*, *SAGE1;* pooled data). Cluster percentages were first analyzed for vehicle-treated iMGLs (**Figure 3A-E**). While most clusters were unaffected, we observed an increased percentage of INPP5D KD cells in cluster 6 (HLA; **Figure 3B**) and a decreased percentage of MS4A6A KD cells in cluster 1 (DAM; **Figure 3D**). Under vehicle conditions, there were no significant population shifts for RABEP1 and EED (**Figure 3C and 3E**). These findings suggest that under steady-state conditions, knockdown of *INPP5D* and *MS4A6A* are sufficient to affect a class of immune signaling and “DAM-like” responses respectively. Chemical inhibition of INPP5D’s phosphatase activity in iMGLs increased inflammasome-related signaling^37^ and MS4A locus members are associated with regulation of soluble TREM2 (sTREM2) levels^38,39^ (see Discussion), which is conceptually in line with our observations. We next analyzed cell percentages for poly(I:C)-treated iMGLs (**Figure 3F-J**). For those targets in our low ROS group (*INPP5D*, *RABEP1*), we observed a depletion of clusters 3 and 5 (**Figure 3G-H**) beyond what was already caused by poly(I:C) treatment, suggesting that these genes may normally help maintain microglia in a homeostatic state or damper their rapid response to inflammatory stimuli. We also saw a slight enrichment of cluster 6 in INPP5D KD iMGLs, (similar to what was seen in steady state conditions). Importantly, the proportion of cluster 2 ((poly(I:C)-driven inflammatory) did not change, suggesting that our low ROS group did not affect the decision to generate an inflammatory response to poly(I:C) when knocked down, but instead might shape that response. For those targets in our high ROX group (*MS4A6A* and *EED*), we observed a depletion of cluster 0 (one of the clusters associated with homeostatic microglia) and enrichment of cells in cluster 2 (poly(I:C)-driven inflammatory; **Figure 3I-J**). This is consistent with both genes driving an inflammatory response when knocked down. For MS4A6A, we observed a modest depletion of cluster 1 (DAM; **Figure 3I**) like what was seen in the vehicle condition, suggesting it may play a role in maintaining the DAM state. For EED KD iMGLs, we observed additional differences in cluster proportions, specifically depletions in clusters 0, 3, 4, and 5 (**Figure 3J**). These data suggest that *EED* may play a more global role than *MS4A6A* in maintaining cells in the homeostatic and cell proliferating state when exposed to poly(I:C). Alternatively, loss of EED may drive inflammatory signaling more strongly than loss of MS4A6A, as evidenced by the larger increase in the proportion cluster 2 in EED KD conditions (**Figure 3**).

**Figure 3:**
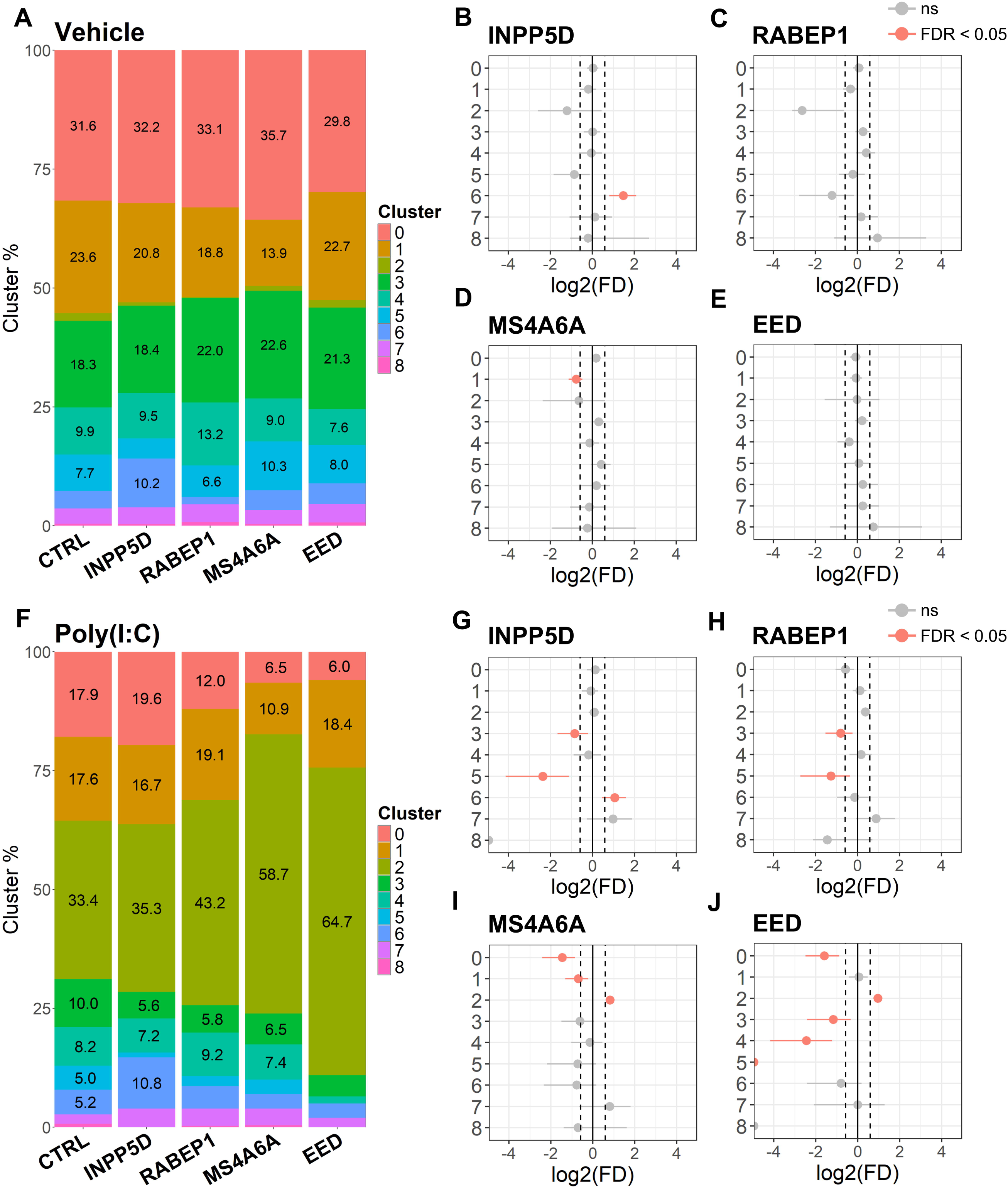
CRISPRi-mediated KD of target genes affects iMGL cluster proportions. (**A**) Cluster percentages for vehicle-treated iMGLs across CRISPRi targets (2,899 total cells). (**B-E**) Point-range plots showing observed log_2_ fold difference for each CRISPRi target in vehicle-treated iMGLs. Dotted lines indicate a minimum log_2_ fold difference of 0.5 and red colored points indicates significance (FDR < 0.05). (**F**) Cluster percentages for poly(I:C)-treated iMGLs across CRISPRi targets for only perturbed cells (2,973 total cells). (**G-J**) Point-range plots showing observed log_2_ fold difference for each CRISPRi target in poly(I:C)-treated iMGLs. Dotted lines indicate a minimum log_2_ fold difference of 0.5 and red colored points indicates significance (FDR).

### 3.7 KD of ROS modulators change global gene expression

Next, we analyzed the transcriptomic changes associated with KD of CRISPRi targets. We compared the downregulated (DR) and upregulated (UR) DEGs between vehicle and poly(I:C)-treated iMGLs for low and high ROS targets, across all clusters (**Figure 4A-D, Table S4**). Interestingly, there were many DEGs in low ROS targets under vehicle conditions (226 shared DR DEG and 220 shared UR DEGs), many of which also overlapped with poly(I:C)-treated iMGLs (**Figure 4A-B**). This was not observed for high ROS targets, where there were little to no DEGs under vehicle conditions (**Figure 4C-D**). However, with poly(I:C) treatment *MS4A6A* KD iMGLs had many DR and UR DEGs, 311 and 524 DEGs, respectively (**Figure 4C-D**), while *EED* had 59 UR DEGs, 22 of which overlapped with *MS4A6A* UR DEGs, and 1 DR DEG. *MS4A6A* and *EED* thus appear to have a greater effect on microglial state when challenged with an inflammatory stimulus than under steady state conditions.

**Figure 4:**
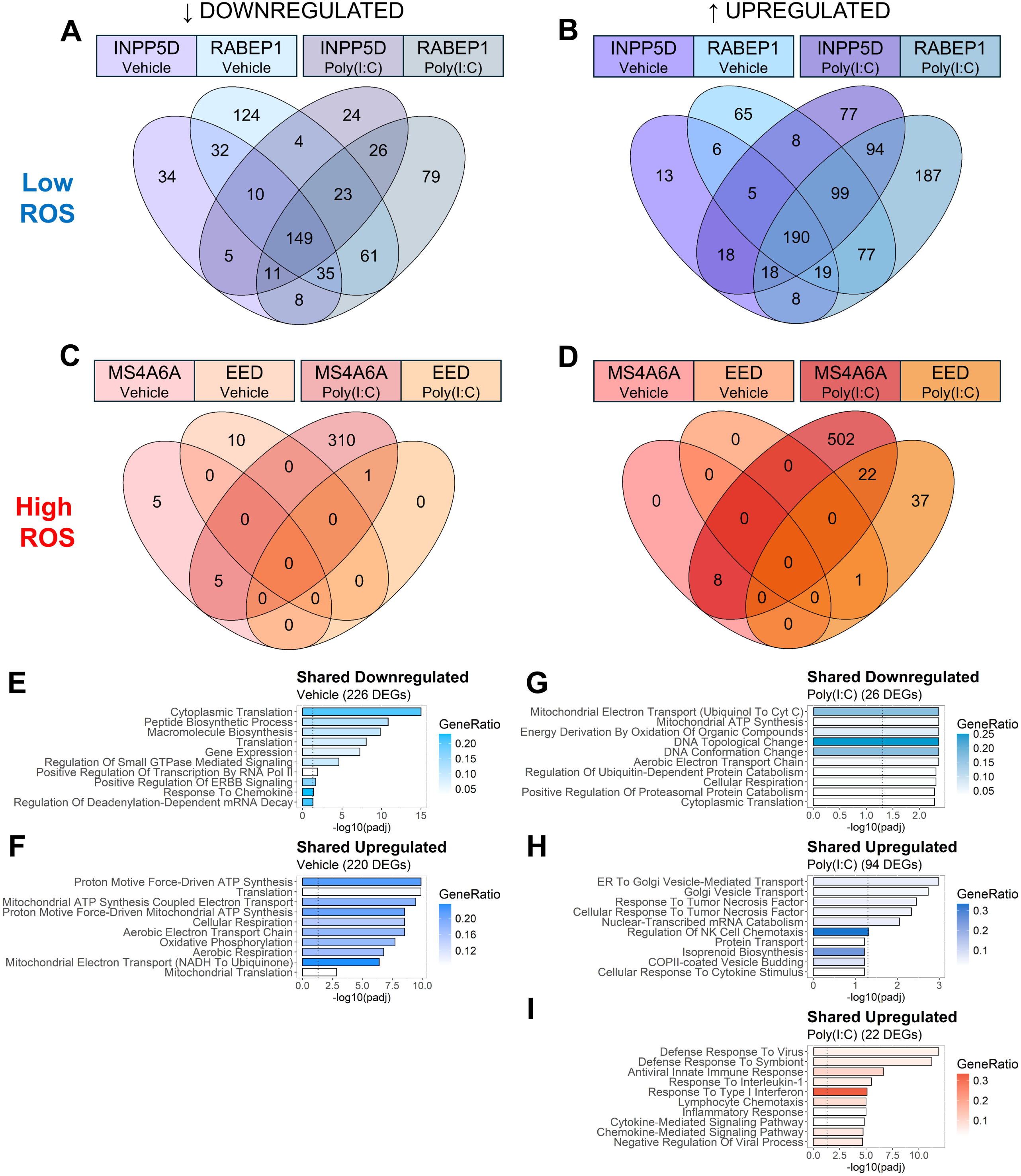
Inhibition of target genes affects iMGL gene expression and pathways. (**A-B**) Venn diagrams comparing vehicle and poly(I:C) transcriptional responses for low ROS CRISPRi targets (*INPP5D* and *RABEP1*, purple and blue, respectively) and (**C-D**) high ROS CRISPRi targets (*MS4A6A* and *EED*, red and orange, respectively). Downregulated and upregulated DEGs were filtered to include those with adjusted p-values ≤ 0.05 and log2 fold change ≥ 0.25. (E-F) Pathway analysis showing significant pathways for downregulated (**E**) and upregulated (**F**) shared vehicle DEGs in low ROS CRISPRi targets. (**G-H**) Pathway analysis showing significant pathways for downregulated (**G**) and upregulated (**H**) shared poly(I:C) DEGs in low ROS CRISPRi targets, excluding changes seen under vehicle conditions. (**I**) Pathway analysis showing significant pathways for upregulated shared poly(I:C) DEGs in high ROS CRISPRi targets, excluding changes seen in vehicle conditions. The top 10 GO biological processes are colored according to the gene ratio, with the dotted line showing - log10(0.05) significance threshold. Significant pathways had adjusted p-values ≤ 0.05.

To better understand how *INPP5D* and *RABE1* KD might similarly affect microglial function, we performed pathway analysis on shared DR and UR DEGs in these low ROS targets under vehicle conditions (**Figure 4E-F and Table S5**). DR pathways were associated with core cellular processes, such as cytoplasmic translation, macromolecular biosynthesis, and gene expression. Additionally, there were also pathways associated with response to cytokines and small GTPase signaling (**Figure 4E**). UR pathways included ATP synthesis, cellular respiration, and oxidative respiration (**Figure 4F**). This suggests that KD of either low ROS gene greatly affects basal cellular metabolism which we hypothesize may influence responses to inflammatory stimuli. In support of this idea, oxidative phosphorylation energy production is associated with homeostatic microglia and anti-inflammatory stimulation, which switches to glycolysis under inflammatory conditions in a manner similar to the Warburg effect in cancer cells^40,41^. Next, we performed pathway analysis on shared DR and UR DEGs in low ROS targets under poly(I:C)-stimulated conditions (**Figure 4G-I and Table S5**). We observed many significant DR pathways, which were associated with ATP synthesis, cellular respiration, and electron transport chain (**Figure 4G**). Conversely, significant UR pathways were associated with Golgi vesicle transport, response to tumor necrosis factor (TNF), and regulation of NK cell chemotaxis (**Figure 4H**). The reduction in oxidative phosphorylation-related pathways is likely an inflammatory shift away from their elevated steady-state levels while in parallel distinct inflammatory-related pathways are activated. To better understand shared *MS4A6A* and *EED* KD responses, we performed pathway analysis on shared UR DEGs in these high ROS targets under poly(I:C)-stimulated conditions (**Figure 4I and Table S5**). We observed several UR pathways associated with defense response to virus, antiviral immune response, and cytokine-mediated signaling (**Figure 4I**). Overall, it appears that KD of high ROS targets leads to a heightened immune response to poly(I:C) stimulus, which is consistent with effects on inflammatory cluster 2 proportion (**Figure 2**).

Finally, we wanted to further investigate pro-inflammatory cluster 2-specific changes as we observed an enrichment of this poly(I:C)-driven inflammatory cluster in KD of high ROS targets. This cluster also has significant overlap with CRM2 (cytokine release microglia) and IRM (interferon release microglial) populations seen in xMGLs in an AD mouse model^1^, and taken together suggests it is the most AD-relevant cluster for damaging inflammatory effects. We compared UR DEGs for each CRISPRi target in poly(I:C)-treated iMGLs only in cluster 2 (**Figure S3A**). Analysis of *INPP5D* unique UR DEGs revealed several pathways associated with MHCII protein complex assembly, antigen presentation, and immunoglobulin production (**Figure S3B**). This is in line with its effects on HLA cluster 6 and suggests a largely unappreciated role for *INPP5D* in regulating these functions. For *RABEP1* unique UR DEGs, we saw pathways associated with macromolecule biosynthesis, mitochondrion organization, and protein stabilization (**Figure S2C**). Potentially these pathways result from changes in early endosomes function associated with RABEP1 protein activity^42–44^, but the precise mechanism and functional consequence are currently unclear. KD of *MS4A6A* was associated with 3 significant UR pathways involved in iron-sulfur cluster assembly (**Figure S3D**). Iron-sulfur clusters in the mitochondria have physiological roles in electron transport in Complexes I-III but also help drive mitochondrial ROS (reviewed in ^45^) which could potentiate inflammatory responses. Lastly, we checked if pro-inflammatory gene expression for genes in cluster 2 were affected by KD of CRISPRi targets. Indeed, we show that pro-inflammatory and antiviral genes had higher expression in iMGLs with *MS4A6A* and *EED* KD (high poly(I:C)-induced ROS when KD; **Figure S3E**). In contrast, *INPP5D* and *RABEP1* KD (low poly(I:C)-induced ROS when KD) had lower expression of these genes (**Figure S3E**).

Overall, our findings suggest that KD of low ROS targets (*INPP5D*, *RABEP1*) have many shared responses under basal conditions and when stimulated with poly(I:C), including similar changes in cellular metabolism, although each also has distinct inflammatory responses. On the other hand, KD of high ROS targets (*MS4A6A*, *EED*) does not lead to many gene expression changes under basal conditions while when challenged with poly(I:C) each leads to upregulation of genes associated with a heightened immune response.

## 4 Discussion

Our goal was to develop a high throughput method to begin to understand how AD GWAS hits affect microglial inflammation in a human-specific model. Our strategy was to develop an assay with a fluorescent readout of an inflammatory process that could be used to identify target genes associated with different levels of inflammatory activation. To this end, we conducted high throughput CRISPRi screens in iMGLs with ROS production as a readout in response to the viral mimic poly(I:C) (**Figure 1**). We chose poly(I:C) as a model since microbial pathogens such as viruses have been associated with AD and Aβ has been hypothesized to increase as a protective response to viruses and other microbes^46–48^. Future studies focusing on additional inflammatory stimuli such oligomeric Aβ42, which we have shown transiently drives inflammatory processes in iMGLs^14^, will be particularly impactful since they will allow us to determine how much of our findings are general versus specific to the particular inflammatory stimulus selected.

119 AD GWAS risk factors^28,29^ were analyzed in parallel for ROS production via CRISPRi modulation, along with negative control non-essential genes and positive control essential genes that would be expected to get depleted in the screen (**Figure S1).** 7 genes were consistently in the top 20 drivers or repressors of ROS (**Figure 1E**). Genes that when knocked down reduced ROS included *INPP5D*, *RABEP1*, and *PVR*, and genes that elevated ROS included *MS4A6A*, *EED*, *AGFG2*, and *CECAM16*. We then further assessed low and high ROS hits by knocking them down individually using CRISPRi and conducting scRNA-seq of vehicle and poly(I:C) treated iMGLs in 2 independent CROP-seq experiments. *INPP5D*, *RABEP1*, *MSA4A6A*, and *EED* had significant effects and were analyzed further.

We identified 9 independent clusters in our iMGLs (**Figure 2**). Poly(I:C) treatment itself changed cluster proportions, including the appearance of a poly(I:C)-driven inflammatory cluster 2, as 97% of the cells in cluster 2 were found in the poly(I:C) condition (**Table S4**). Importantly, marker genes in this cluster have been observed in xMGLs in a humanized AD mouse model^1^. They identified 2 cytokine response clusters which have overlapping genes with those found in cluster 2, suggesting that poly(I:C) cytokine responses may also reflect human responses to AD pathology. Several other clusters are reduced by poly(I:C), including clusters 0, 3 and 5. Cluster 0 is the largest cluster under steady-state conditions (31% of cells) and probably represents a homeostatic population as other scRNA-seq of human microglia have reported^9^. Clusters 3 and 5 are less defined, although they share properties with previously described homeostatic microglia. While cluster 3 has no defined pathways for upregulated genes, cluster 5 is associated with G-protein coupled receptor signaling and astrocyte activation amongst other GO terms (**Table S4**). Further studies will be needed to better understand the functional roles for cluster 3 and 5 cells and their relation to cluster 0.

CRISPRi modulation of low and high ROS targets changed the proportion of cell within each cluster under steady-state and poly(I:C) conditions (**Figure 3**). KD of *MS4A6A* and *EED*, the high ROS hits, led to increased cluster 2 (poly(I:C)-specific antiviral inflammatory) and reduction of cluster 0 (homeostatic). *MS4A6A* KD led to reduction of cluster 1 (DAM) in both vehicle and poly(I:C) treatment conditions, suggesting that the DAM-like response in human microglia might be impaired by LOF in this gene. Although few DEGs are present under vehicle conditions from *MS4A6A* KD (**Figure 4),** DR DEGs during poly(I:C) treatment include multiple transcription factors associated with the DLAM (DAM and lipid associated macrophages) response (*BHLHE41*, *ID2*, *MAF*, *MEF2A*, *MEF2C*) which could play a part in this diminished response^49^. *EED* KD also led to reduction of cluster 3 and 5, most likely just augmenting these reductions that also occur by poly(I:C) treatment, and the proliferative cluster 4. On the other hand, KD of low ROS genes *INPP5D* and *RABEP1* did not lower cluster 2 proportions but did lead to a large number of differentially expressed genes, suggesting that LOF of these genes does not affect the initiation of an inflammatory response, but rather shape that response (**Figure S2**). Similar to *EED* KD, *INPP5D* and *RABEP1* KD also reduced clusters 3 and 5 which are associated with the homeostatic state and may suggest both genes play roles in maintenance of a quiescent state under unperturbed conditions. Finally, *INPP5D* KD under either vehicle or poly(I:C) conditions increased poly(I:C)-independent immune cluster 6, suggesting that’s *INPP5D*’s role in inflammatory processes is more complex.

Our data suggests that LOF of our high ROS AD GWAS hits *EED* and *MS4A6A* drives inflammatory processes in microglia. For *EED* (*<u>E</u>mbryonic <u>E</u>ctoderm <u>D</u>evelopment*), a transcription-wide association study (*TWAS*) suggested that lower expression of *EED* would be deleterious for AD risk^50^. A mouse microglial-specific *EED* knockout model showed altered microglial morphology and augmented synapse pruning that led to learning and memory deficits^51^, demonstrating that loss of EED led to overly activated microglia consistent with our data. While the mechanisms of *EED’s* effects on microglia is unclear, it is a key component of the Polycomb Repressor Complex 2 (PRC2) which leads to histone H3 methylation at lysine 27 to transcriptionally silence genes^52^. Mutations in *EED* can cause Cohen-Gibson Syndrome, one of several PRC2 overgrowth disorders with different PRC2 components affected which are associated with intellectual disability in addition to overgrowth and skeletal abnormalities^53^. Our data supports the idea that altered microglial activity could play a pathogenic role in the neurodevelopmental component of PRC2 overgrowth disorders. Future studies should address how *EED* KD affects human microglial chromatin states to elucidate if derepression of inflammatory genes is the mechanism how its KD increases inflammation. It should also be noted that our *EED* KD was modest, likely blunting our results, and that other LOF methodologies (i.e., CRISPR KO) should be used in the future.

The AD literature for *MS4A6A* (*Membrane Spanning 4-Domains A6A*) and the entire *MS4A* locus containing 13 gene members is more complicated. Multiple family *MS4A* family members have been associated with AD^28,29,54^, including *MS4A2*, *MS4A3*, *MS4A4A*, *MS4A4E*, *MS4A6A*, and *MS4A6E*, all of which were tested in our CRISPRi screens. Interestingly, only *MS4A6A* appeared consistently between both screens as a top 20 high ROS hit (**Table S1**). There is a protective SNP in *MS4A6A* (rs610932) for AD^55^ which is associated with lower expression in blood^56^ and decreased atrophy for particular left-brain structures in AD patients^57^, which would argue that lower expression of *MSA4A6A* would be protective. Several studies suggest MS4A locus genes can modulate soluble TREM2 (sTREM2) levels in the CSF, albeit with contradicting claims. sTREM2 is generated either by ADAM-protease cleavage of full length TREM2 or by alternative splicing and is correlated with AD progression although generally it is thought to be protective^58^. A recent preprint argued MS4A4A protein acts by stabilizing MS4A6A protein which in turn binds TREM2 and inhibits its cleavage into sTREM2 (and ICD)^39^. This was preclinical data for a monoclonal antibody currently in clinical trials to inhibit *MS4A4A* to increase TREM2 signaling. On the other hand, another study showed knockdown of *MS4A4A* or targeting it with an antibody decreased sTREM2 levels whereas overexpression of MS4A4A increased sTREM2 levels, showing a positive relationship between MS4A4A and TREM2 signaling and arguing for compounds that promote MS4A4A levels as a therapeutic approach^38^. This study also showed a missense mutation in *MS4A6A* (pA112T) was associated with higher sTREM2 levels and decreased AD risk, although whether this was a LOF or GOF mutation is currently unclear. Interestingly, our data showed that *MS4A6A* knockdown negatively affected the proportion of iMGLs in the DAM cluster 1 (**Figure 2**), consistent with *MS4A6A* affecting TREM2 signaling. Future studies will need to determine the functional consequences of those DAM-related transcriptional changes and further elucidate the mechanistic relationships between different *MS4A* locus members.

Our low ROS hits (*INPP5D*, *RABEP1*) show a modulation of inflammatory signaling and significant gene expression changes in both steady state (vehicle) and poly(I:C) conditions (**Figures 4, S3**). *INPP5D* encodes src homology 2 (SH2) domain-containing inositol 5’ phosphatase 1 (SHIP1), which converts PI(3,4,5)P_3_ into PI(3,4)P_2_ to negatively regulate multiple signal cascades including phosphoinositide 3-kinase (PI3K)^59^. *INPP5D* has been repeatedly identified as a risk gene involved in AD with both risk and protective variants^28,60,61^, with these SNPs being significantly associated with CERAD score^62,63^. Analysis of human AD samples have shown that AD subjects have increased INPP5D expression^60,64^, although one recent study using antibodies against different epitopes of INPP5D protein reported loss of full length INPP5D but increased in truncated INPP5D^37^. Some AD mouse models support benefits from reduced INPP5D. For instance, *INNP5D* haploinsufficiency in 5xFAD mice led to reduced diffuse plaques, increased dense-core plaques, and improved working memory^65^. However, conditional microglia-specific knockout in an APP^Swe^/PSEN1^ΔE9^ mouse model showed increased plaques^66^, albeit without negative effects on synapses, suggesting contextual differences of *INPP5D* LOF. INPP5D has also been shown to interact with TYROBP (DAP12, a direct downstream mediator of TREM2 signaling), leading to inhibition of cellular activation^67,68^. Studies in the humanized mouse model *APP^NL-G-F^* with *TYROBP* KO and *INNP5D* haploinsufficiency showed that reduced *INNP5D* rescued microglial localization around Aβ plaques, improved plaque compaction, and reduced dystrophic neurites observed in *TYROBP* KO mice^69^. The precise role of *INPP5D* in human microglia in the context of AD is also not clear. In a recent study iMGLs were treated with a selective INPP5D inhibitor and showed activation of the inflammasome and increased secretion of pro-inflammatory IL-1β and IL18^37^, which could have negative consequences for AD progression. While in our data *INPP5D* KD produced lower ROS in response to poly(I:C), it also increased HLA cluster 6 under steady-state and poly(I:C) conditions (**Figure 3B,G**). The functional consequences of modulating this HLA/poly(I:C)-independent inflammatory cluster are currently unclear. As variants of HLA genes are associated with increased AD risk^28,29,70^, is essential to better understand the functions of HLA-related genes in microglia in the context of AD and how they react with recruited immune cells from the periphery and other CNS-resident cells. Further studies could also address the role of specific *INPP5D* risk variants, whether its phosphatase activity is required for all INPP5D microglial functions in the context of AD, and whether INPP5D may play different roles in different stages of AD.

Although *RABEP1* knockdown had significant overlap with *INPP5D* knockdown in terms of DEGs (**Figure 4**), RABEP1 acts by a completely different mechanism and to our knowledge has no functional link with INPP5D outside of existing AD GWAS studies. RABEP1 regulates endocytic trafficking, including membrane docking, fusion, and recycling vesicles^42–44^. In addition to non-coding SNPs ^29,54,71^, a missense variant p.R845W was identified as being associated with AD in a Chinese cohort. When that variant was knocked into HEK 293 cells, it led to decreased proliferation, enlarged early-endosomes, and elevated Aβ42:40 ratio^71^. Interestingly, a SNP associated with longevity in a long-lived family was found in the 3’ UTR of *RABEP1*, although it also created a missense mutation in *NUP88* on the other strand, so which gene is responsible (or both) for potential longevity effects is unclear^72^. How presumed defects in endocytic trafficking led to decreased microglial inflammation and ROS production by *RABEP1* knockdown in iMGLs remains unclear, although one possibility is it changes poly(I:C) endocytosis and interaction with the poly(I:C) receptor TLR3 in the ER. However, this poly(I:C) endocytic mechanism would not explain the strong effect of RABEP1 knockdown on steady-state microglial transcriptome (**Figure 4**), and why this overlaps significantly with *INPP5D* knockdown, suggesting that additional investigation is required to understand its underlying mechanism. Our data showing these low ROS hits overlap on effects on cellular metabolism may provide a framework to begin to answer these questions.

There are several limitations to our study. First, our CRISPRi screen was only focused on ROS production as our microglial inflammatory output, which does not reflect the full repertoire of inflammatory responses such as secretion of cytokines and chemokines. However, for the subset of genes explored via CROP-seq, we obtained a more global view of their effects including ones on cytokine and chemokine production. Furthermore, the initial screening pipeline developed here could be successfully applied to other inflammatory features such as specific cytokines or signal transduction pathways. Secondly, we did not look at any effects of CRIPSRi of target genes on non-inflammatory processes such as phagocytosis, which should be assessed in the future. Finally, all experiments were conducted *in vitro*. Xenotransplantation of CRISPR manipulated iMGLs into control or AD mouse models in future studies would enable further maturation of iMGLs and studies of interactions with host tissue.

## Supporting information

Supplemental Table Inventory

## Acknowledgements

We would like to thank Paroma Mallick for her assistance with the design of our CRISPRi library.

## Author Contributions

CLC, LW, JW, EA, GS, RP, NI conducted experiments or did substantial analysis of data. SC generated tools used for experiments. CLC, AC and AAS drafted the manuscript with assistance from all the authors. AFT and PC provided significant advice and technical insight on the project. AAS, AC, and HG were responsible for conceptualization and management of the project. All authors edited the manuscript.

## Notes

**Funding statement:** This work was primarily supported by a Development Grant (A.C., A.A.S., G.H. Co-PIs) from the Columbia University Alzheimer’s Disease Research Center (ADRC) associated with P30AG066462-02. A.A.S. is also supported by NIA RF1AG078352, The Henry and Marilyn Taub Foundation, and The Thompson Family Foundation (TAME-AD). A.C. is supported by a Career Awards for Medical Scientists from the Burroughs Wellcome Fund, a DP2 award (DP2NS131566-02) from the NIH, and a Collaborative Pairs grant from the CZI. C.L.C. is supported by a Diversity Fellowship associated with R01AG073360 (A.F.T. PI).

### Summary of Updates

This version changes terminology and includes an additional reference.

