## Supplemental Figure S1 for "High throughput identification of genetic regulators of microglial inflammatory processes in Alzheimer’s disease"

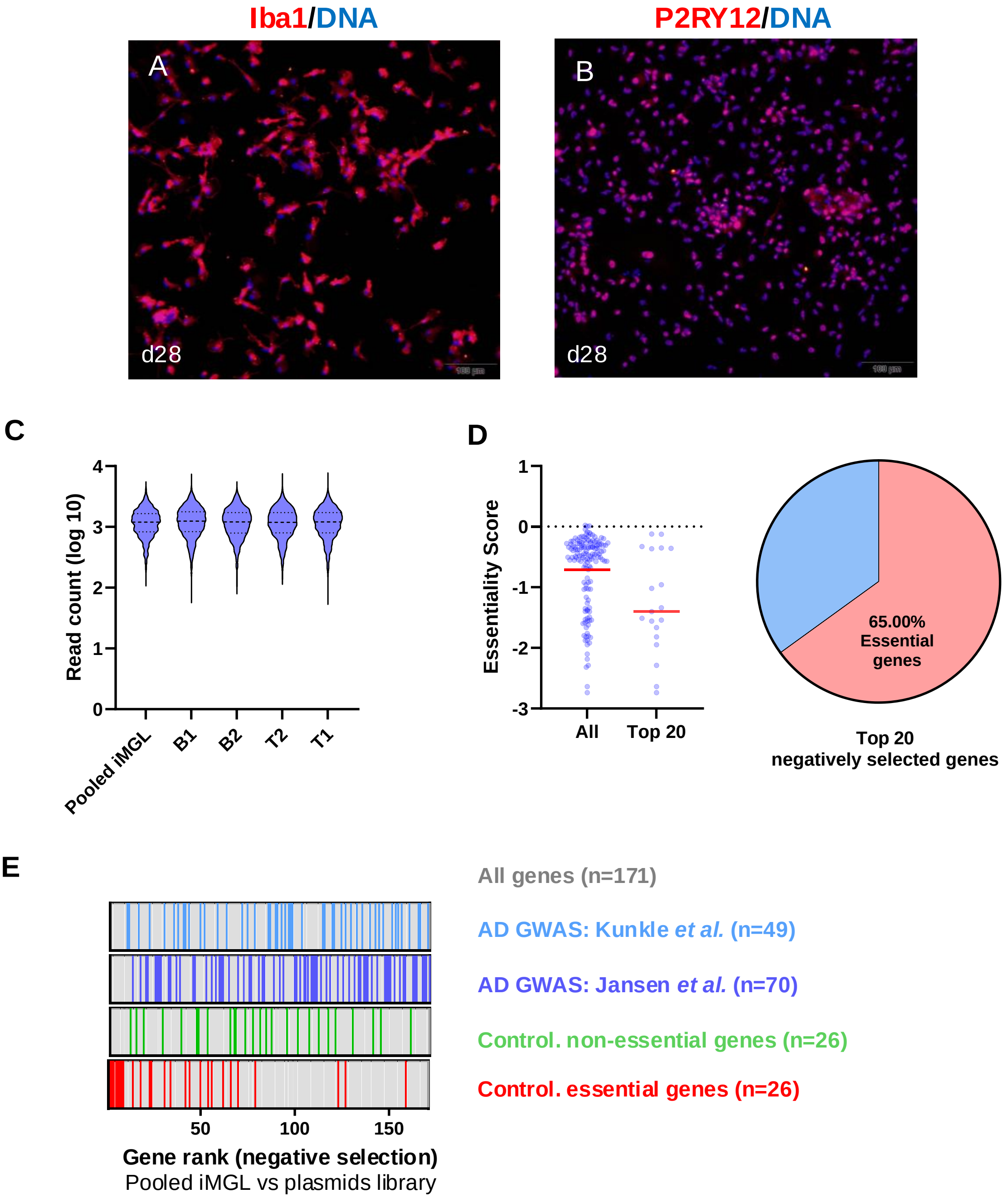


**Figure S1. Quality control analysis of differentiation and CRISPR-Cas9 screen in induced microglia-like cells (iMGLs). (**A-B**)** Immunostaining of iMGLs showed most cells expressed the microglial markers Iba1 (A; red) and P2RY12 (B; red). Nuclei are stained with DAPI (blue) and the scale bar = 100 μM. (C) Distribution of guide RNA (gRNA) normalized read counts across pooled iMGLs and all sorted bins in the CRISPR-Cas9 screen. The data demonstrate consistent gRNA representation and coverage across sorted populations with medium read count around 1000, validating the library complexity and sequencing depth. (D) Median essentiality scores of the top 20 negatively selected genes from the screen. Lower essentiality scores indicate higher essentiality, which demonstrated a depletion effect, validating the effectiveness of the screening process. (E) Distribution of genes across different categories in the negative selection results of the screen. Essential genes are significantly enriched on the negative selection side, whereas non-essential genes and genes derived from Kunkle and Jansen's studies are not enriched on either side. These results validate the consistency and effectiveness of the CRISPR-Cas9 screening approach in iMGLs.
