## Supplementary figures and images for "High throughput identification of genetic regulators of microglial inflammatory processes in Alzheimer’s disease"

### Supplemental Figure S2

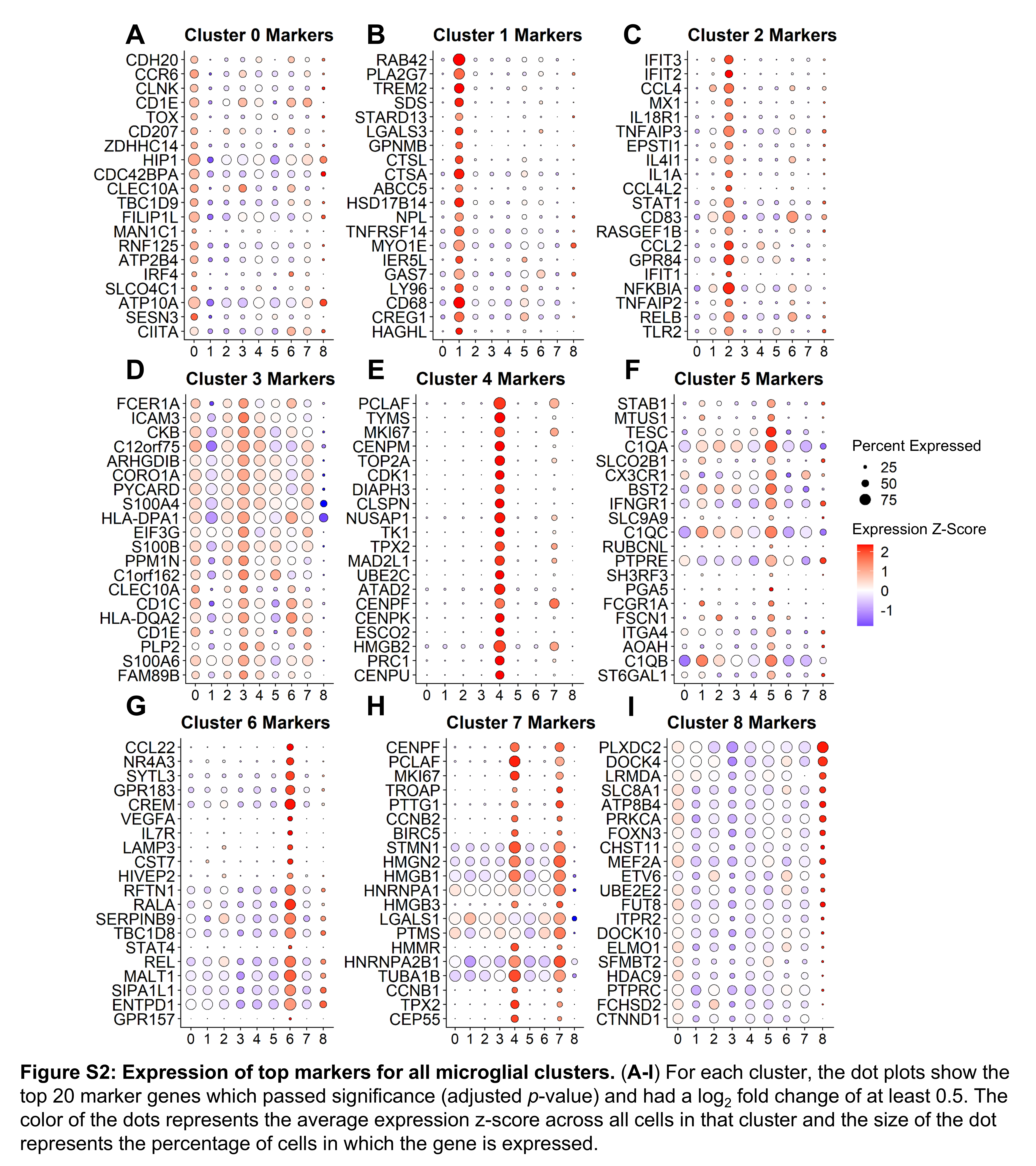

### Supplemental Figure S3

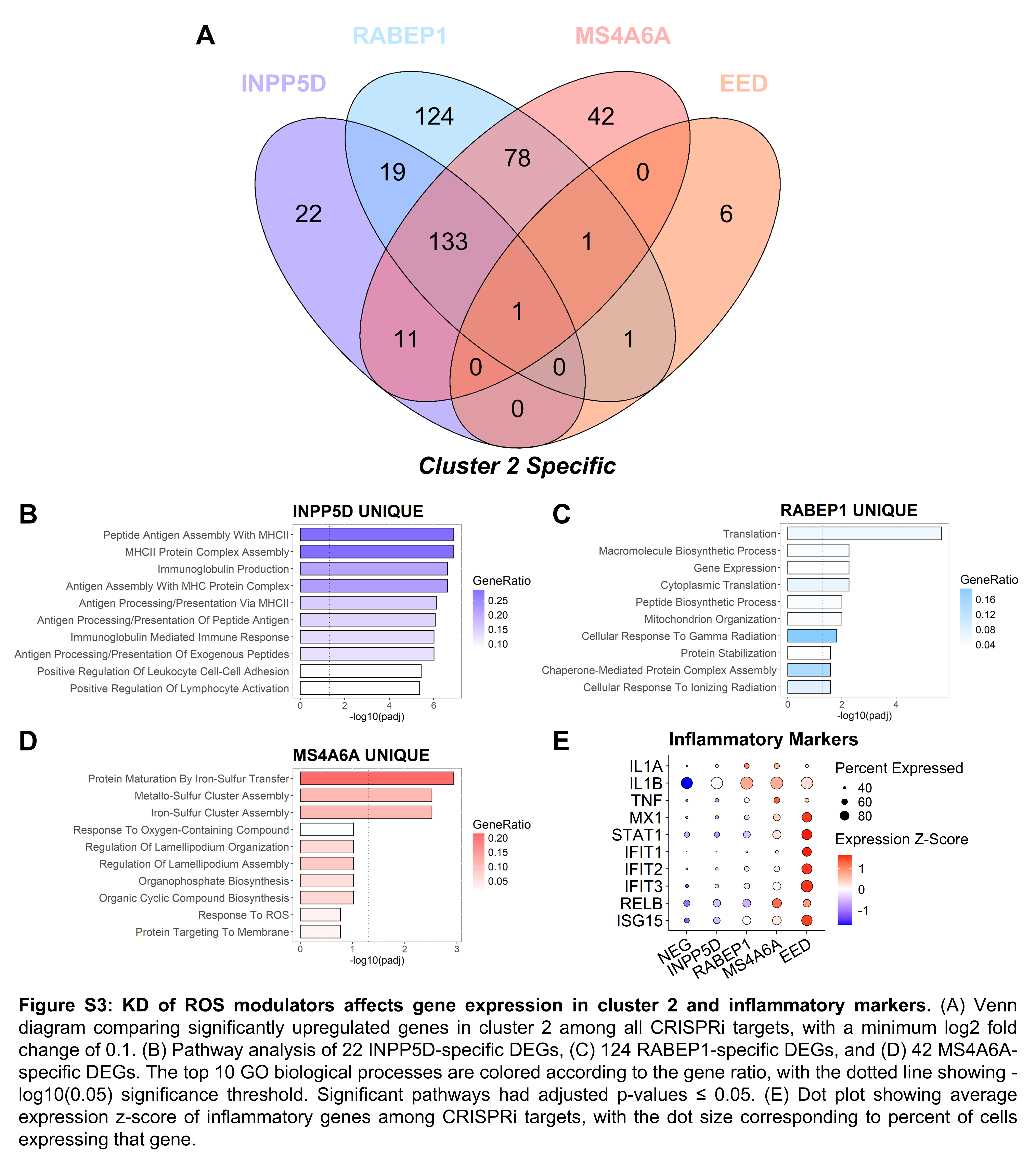
