## Supplemental Table Inventory for "High throughput identification of genetic regulators of microglial inflammatory processes in Alzheimer’s disease"

**Table S1: Genes in CRISPRi library.** This table relates to Figure 1. It includes all gRNAs used in the study, including breakdown into essential gRNAs, non-essential gRNAs, Kunkle et. al GWAS target gRNAs, and Jansen et. al GWAS target gRNAs. It also shows the top 20 negative and positive hits from 2 independent CRISPRi screens.

**Table S2: gRNAs used Peturb-seq experiments.** This table is part of the Methods and relates to Figures 2-4. It includes gRNA sequences for the 8 genes targeted for Peturb-seq experiments, including negative controls *CABP5* and *SAGE1*.

**Table S3**: **Top 25 depleted genes depleted by CRISPRi.** This Table relates to Figure 1, Figure S1. It shows the top 25 genes depleted in CRISPRi screens and the associated DepMap score to see if it would be predicted to be essential.

**Table S4: Analysis and characterization of iMGL clusters.** This table relates to Figure 2. Tab 1 shows cluster % by treatment type. Tabs 2-3 the DEGs when comparing iMGLs treated with poly(I:C) compared to vehicle controls and the pathways associated with those DEGs. The cluster markers tab shows significant markers for all clusters. The tabs labeled cluster 0 through 8 shows the full pathway analysis results from enrichR. The final module scores tab reports the average module scores across clusters for the top genes in all clusters from the Mancuso et al. study. The tab also shows the module score z-score with their adjusted p-value across all clusters.

**Table S5: Pathway analysis comparing KD of ROS regulators.** This table relates to Figure 4. These tabs show the full results from pathway analysis using enrichR of groups identified in Figure 4.
