## Supplemental Table S2 for "High throughput identification of genetic regulators of microglial inflammatory processes in Alzheimer’s disease"

**Table S2. Spacer sequences used within Perturb-seq experiments.**

| Gene | Spacer sequence |
| --- | --- |
| AGFG2 | GGAAGGCGGCAGTGGTTGAA |
|  | CAGGGGAGCCGGGCGTGCGG |
|  | GAAGGCGGCAGTGGTTGAAG |
| EED | AAACGTCTTTGGAAGGAGGA |
|  | AACGTCTTTGGAAGGAGGAA |
|  | GCGGCTGAAACGTCTTTGGA |
| MS4A6A | AGTCCTCAGAAGCTCCAAAG |
|  | AACCAGAGTTAAAACCTCTT |
|  | TGGTTTCTCAGTCCCATCAA |
| INPP5D | GGCGGCAGGTTGCAGTGGAG |
|  | GGCCACCAAGAGGCAACGGG |
|  | CAACGGGCGGCAGGTTGCAG |
| RABEP1 | GGAGGCGGAGGTCGGCGGTC |
|  | GTCTCTGCCCGCGGCTGTGG |
|  | GCCGGCGCCGCCACAGCCGC |
| PVR | CAGGTCCTGGAATCCCCGGG |
|  | ACTGGAGGAGCGGCCCCCCG |
|  | CTCAGGTCCTGGAATCCCCG |
| CABP5 | AAGAGGTGGCAAAGGAGTGC |
|  | AAGGAGAAGCCAAGAGAGGC |
|  | CCACCTCTTCCTGCCTCTCT |
| SAGE1 | GCGGGATAAAAGGGTGAATT |
|  | TGCACAGCGGGCAGAGGCAG |
|  | CTTCACTATCAACTGCACAG |
