## Supplemental Table S3 for "High throughput identification of genetic regulators of microglial inflammatory processes in Alzheimer’s disease"

**Table S3. Top 25 depleted genes in pooled CRISPRi library transduced iMGLS**

| **id** | **neg\|score** | **neg\|p-value** | **neg\|fdr** | **neg\|rank** | **neg\|lfc** | **Depmap Essentiality score** | **Marked as Essential in library** |
| --- | --- | --- | --- | --- | --- | --- | --- |
| SPI1 | 3.26E-09 | 4.99E-06 | 0.000427 | 1 | -1.9642 | -1.02 | TRUE |
| POLA1 | 8.83E-07 | 4.99E-06 | 0.000427 | 2 | -1.2635 | -1.54 | TRUE |
| PSMC2 | 2.21E-06 | 1.50E-05 | 0.000853 | 3 | -1.2712 | -1.513 | TRUE |
| RPL9 | 1.14E-05 | 3.49E-05 | 0.001493 | 4 | -1.254 | -2.289 | TRUE |
| NUP98 | 1.41E-05 | 4.49E-05 | 0.001536 | 5 | -0.70865 | -1.403 | TRUE |
| PSMD7 | 1.82E-05 | 5.49E-05 | 0.001564 | 6 | -1.091 | -2.738 | TRUE |
| RPS24 | 0.00026708 | 0.0011327 | 0.027669 | 7 | -0.93252 | -1.818 | TRUE |
| COPA | 0.00049108 | 0.0021306 | 0.045542 | 8 | -0.82866 | -2.185 | TRUE |
| RPS27 | 0.00060114 | 0.0026595 | 0.050531 | 9 | -0.53521 | -1.337 | TRUE |
| INPP5D | 0.00078458 | 0.003398 | 0.058106 | 10 | -0.61215 | -0.107 | FALSE |
| PSMC4 | 0.0020114 | 0.0079187 | 0.117819 | 11 | -0.80724 | -1.164 | TRUE |
| PSMD11 | 0.00213 | 0.008268 | 0.117819 | 12 | -0.74848 | -1.666 | TRUE |
| PSMA3 | 0.0053194 | 0.018487 | 0.243174 | 13 | -0.66565 | -2.639 | TRUE |
| RPL6 | 0.0062643 | 0.021151 | 0.25835 | 14 | -0.59836 | -1.597 | TRUE |
| COPZ1 | 0.0093777 | 0.030043 | 0.342491 | 15 | -0.51753 | -1.559 | TRUE |
| PSMC1 | 0.011537 | 0.036729 | 0.392545 | 16 | -0.20825 | -1.492 | TRUE |
| EPHB4 | 0.014491 | 0.045441 | 0.457087 | 17 | -0.38977 | -0.126 | FALSE |
| PSMC3 | 0.015593 | 0.048455 | 0.460324 | 18 | -0.43323 | -1.88 | TRUE |
| NUP133 | 0.021936 | 0.062057 | 0.538802 | 19 | -0.355 | -1.799 | TRUE |
| PSMD6 | 0.024057 | 0.066159 | 0.538802 | 20 | -0.34239 | -2.103 | TRUE |
| CLASRP | 0.024066 | 0.066169 | 0.538802 | 21 | -0.34775 | -0.96 | TRUE |
| APP | 0.040083 | 0.097943 | 0.761286 | 22 | -0.21506 | -0.328 | FALSE |
| NUP93 | 0.043088 | 0.10358 | 0.770107 | 23 | -0.44094 | -1.876 | TRUE |
| RPL18A | 0.05289 | 0.12197 | 0.846318 | 24 | -0.00502 | -1.761 | TRUE |
| PSMB2 | 0.054574 | 0.12536 | 0.846318 | 25 | -0.33609 | -1.828 | TRUE |
